# Population genomics reveal genetic variants associated with lunar-regulated spawning time in grass puffer

**DOI:** 10.64898/2026.03.31.715739

**Authors:** Yuma Katada, Daisuke Kurokawa, Mats E. Pettersson, Junfeng Chen, Liang Ren, Taiki Yamaguchi, Tomoya Nakayama, Kousuke Okimura, Michiyo Maruyama, Ren Enomoto, Hironori Ando, Gen Kurosawa, Asako Sugimura, Yoko Hattori, Leif Andersson, Takashi Yoshimura

**Affiliations:** Institute of Transformative Bio-Molecules (WPI-ITbM), Nagoya University, Furo-cho, Chikusa-ku, Nagoya, Aichi, 464-8601, Japan; Laboratory of Animal Integrative Physiology, Graduate School of Bioagricultural Sciences, Nagoya University, Furo-cho, Chikusa-ku, Nagoya, Aichi, 464-8601, Japan; Misaki Marine Biological Station, Graduate School of Science, The University of Tokyo, 1024 Koajiro, Misaki, Miura, Kanagawa 238-0225, Japan; Science for Life Laboratory, Department of Medical Biochemistry and Microbiology, Uppsala University, Uppsala, Sweden; Sado Marine Biological Station, Sado Island Center for Ecological Sustainability, Niigata University, 87 Tassha, Sado 952-2135, Niigata, Japan; RIKEN Center for Interdisciplinary Theoretical and Mathematical Sciences (iTHEMS), Wako, Saitama, 351-0198 Japan; Toyota Boshoku Corporation, 1-1 Toyoda-cho, Kariya, Aichi, 448-8651, Japan; Department of Veterinary Integrative Biosciences, Texas A&M University, College Station, USA; Center for One Medicine Innovative Translational Research (COMIT), Nagoya University, Nagoya, Aichi 464-8601, Japan

**Keywords:** geographic variation, grass puffer, semilunar rhythm, circadian rhythm, population genomics, triple CRISPR

## Abstract

High and low tides occur twice a day (every ∼12.4 hours), with the largest tidal ranges during spring tides around new and full moons (every ∼14.765 days). Although lunar cycles shape diverse animal phenotypes, particularly the reproduction of coastal animals, the genetic basis of lunar-related rhythms remains unclear. Since phenotypic variation is a valuable resource for elucidating such mechanisms, we examined geographic variation in the lunar-regulated mass spawning of the grass puffer (*Takifugu alboplumbeus*) along the Japanese coast. Western populations spawned during the first half of the spring tides, whereas eastern populations spawned during the second half. Furthermore, the spawning was restricted to a narrow daily time window that occurred earlier in the western populations than in the eastern ones. Assuming that spawning is regulated by circadian gating, this geographic shift in the daily spawning window could account for the population difference in semilunar spawning timing. Consistent with this hypothesis, the free-running circadian period (τ), which can affect the phase of the gating window, was shorter in the western populations. Mathematical modeling recapitulated the observed geographic difference in the spawning time window based on variation in τ. To identify candidate genes responsible for these differences, we conducted population genomics analysis and identified proline-rich transmembrane protein 1-like (prrt1l) as a candidate. Expression of *prrt1l* was observed in the circadian pacemaker suprachiasmatic nucleus, and triple CRISPR F0 knockout of *prrt1l* shortened the free-running period in larvae. These findings suggest a potential mechanism underlying the geographic variation in lunar-synchronized spawning behavior.

**Highlights:**

- The geographic variation exists in the lunar-regulated spawning of the grass puffer, with differences in spawning dates and times between western and eastern Japan.
- The free-running period of western populations is shorter than that of eastern populations, which is consistent with their earlier spawning timing.
- Population genomics analysis identified *prrt1l* as a candidate gene harboring population-specific missense mutations, the knockout of which shortens the free-running period.

## Introduction

Various physiological functions and behaviors exhibit rhythmic fluctuations, which are regulated by endogenous biological clocks and exogenous environmental factors. One such factor is the lunar cycle, which generates periodic changes in moonlight and tides that influence life phenomena, particularly the reproduction and feeding of coastal organisms^1^. While these phenomena are more common in marine organisms, lunar-synchronized behaviors have also been observed in terrestrial species, including the timing of wildebeest conceptions and badger reproductive behavior, as well as human traits such as sleep duration and menstrual cycles^2–5^. However, the underlying genetic basis of lunar rhythms remains unclear.

Variation in rhythmic phenotypes is a valuable target for exploring the genetic basis of biological clocks. For example, in circadian biology, genetic analyses of individuals with altered circadian phenotypes have led to the identification of circadian-related genes and their functional domains. This has advanced our understanding of circadian clock mechanisms^6–8^. In lunar-related rhythm research, the emergence timing of the marine midge *Clunio marinus*, which emerges at low tide according to a genetically determined circadian and circalunar clocks, has been used as a model to study the geographic variation. Quantitative trait loci analysis of the midge revealed alternative splicing of calcium/calmodulin-dependent protein kinase II, suggesting a mechanism underlying natural adaptation in circadian timing^9^.

In our previous study, we focused on the grass puffer (*Takifugu alboplumbeus*), which exhibits mass spawning around the new and full moons (i.e., during spring tides). We demonstrated that prostaglandin E_2_ is involved in the synchronization of the semilunar mass spawning behavior^10^. Interestingly, the timing of the grass puffer spawning has been suggested to differ between populations in western and eastern Japan^10–14^. Therefore, we investigated the geographic variation in the semilunar spawning rhythms of the grass puffer in order to understand the genetic basis of the semilunar rhythm. First, we characterized geographic variation in the semilunar spawning rhythms using previously published records and our own field observations. We then performed population genomics analysis to identify potential candidate genes. Finally, we used the triple CRISPR F0 knockout method to analyze the functional significance of the candidate gene.

## Results

### Geographic variation in spawning timing between the western and eastern populations

First, we focused on four wild grass puffer populations in Japan: Aburatsubo, Kominato, Minamichita, and Toba (Fig. 1A). We replotted previously published spawning data for the eastern populations (Aburatsubo and Kominato) (Fig. 1B). Although these spawning data were reported decades ago^11,14^, our field surveys in 2020 and 2021 confirmed that these populations still exhibit the same spawning timing. For the Minamichita and Toba populations, we collected field observation data on spawning timing (Fig. 1B). A comparison of spawning dates revealed a distinct geographic variation in spawning: the western populations (Minamichita and Toba) spawned during the first half of the spring tide (i.e., 2-4 days before the new or full moon), while the eastern populations (Aburatsubo and Kominato) spawned during the latter half (4-5 days after the new or full moon) (Fig. 1B). The spawning days of the Minamichita and Toba populations were consistent with a previous study of the population in Murozumi, located further west (Fig. 1A)^12,13^. These results clearly demonstrate the geographic difference in spawning days between western and eastern Japan.

**Fig. 1.**
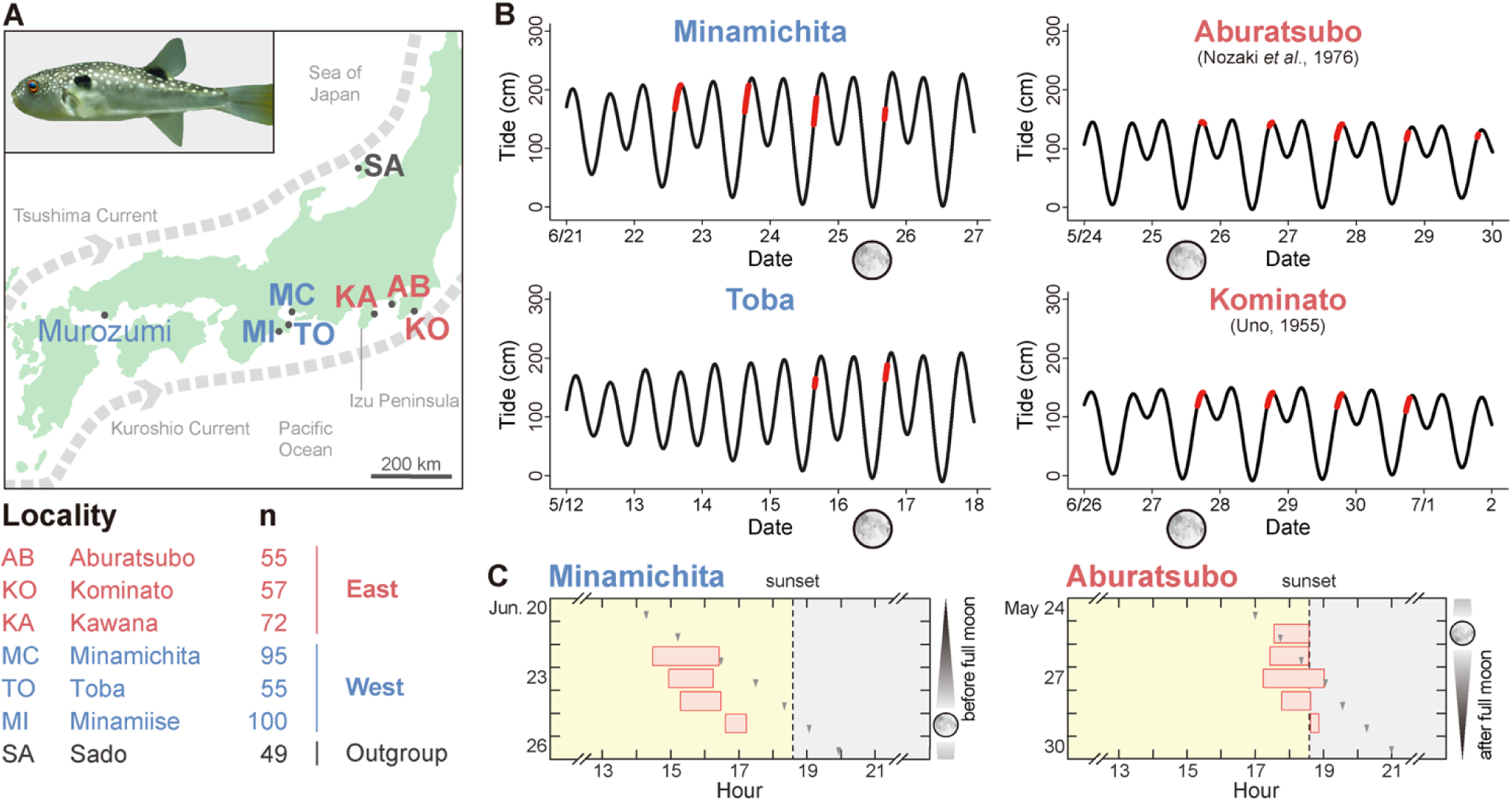
Geographic variation in semilunar spawning of grass puffer. (A) The geographic location of the spawning populations and the number of individuals used in population genomics are shown. (B) The timing of the grass puffer spawning in western and eastern populations. The black and red lines indicate changes in tidal levels and the times at which spawning was observed, respectively. Moon symbols indicate the day of the full moon. The spawning data for Aburatsubo and Kominato were adopted from Nozaki et al. (1976) and Uno (1955), respectively^11,14^. (C) Detailed spawning times of the grass puffer in each population. The red bars indicate spawning times. The grey arrowheads indicate high tides. Moon symbols indicate the day of the full moon. The spawning data for Aburatsubo was adopted from Nozaki et al. (1976)^14^.

Upon closer inspection of the spawning times, we noticed that the spawning times of the day are also different between the western and eastern populations (Fig. 1C). Although onset of spawning was a few hours before high tide in both populations, the western population (Minamichita) spawned around 15:00 to 17:00, whereas the eastern population (Aburatsubo) spawned around 17:00 to 19:00 (Fig. 1C). These results suggest the presence of a population-specific spawning time window, with western populations exhibiting an earlier circadian phase than eastern populations.

### Circadian gating links the spawning time window to the phase of the semilunar spawning rhythm

A circadian gating mechanism provides a mechanistic link between the phase of the spawning time window and the geographic difference in the phase of the semilunar rhythm. Circadian gating restricts biological events to specific time windows defined by the circadian clock^15^. Under this model, spawning occurs only when the potential onset of spawning, which occurs a few hours before high tide in grass puffer, coincides with the circadian controlled time window (Fig. 2A). Although this time window remains fixed at a specific time of day, the potential onset of spawning shift later each day as high tide delays. Thus, the interaction between these two temporal cues can determine the timing of spawning events and, consequently, the phase of the semilunar spawning rhythm (Fig. 2A). For example, if the spawning time window for Minamichita is replaced with that observed in Aburatsubo (eastern Japan), the predicted spawning shifts to the latter half of the spring tide (Fig. 2B). Such simulation demonstrate how the phase of the circadian-controlled spawning window contributes to geographic variation in the semilunar spawning rhythm.

**Fig. 2.**
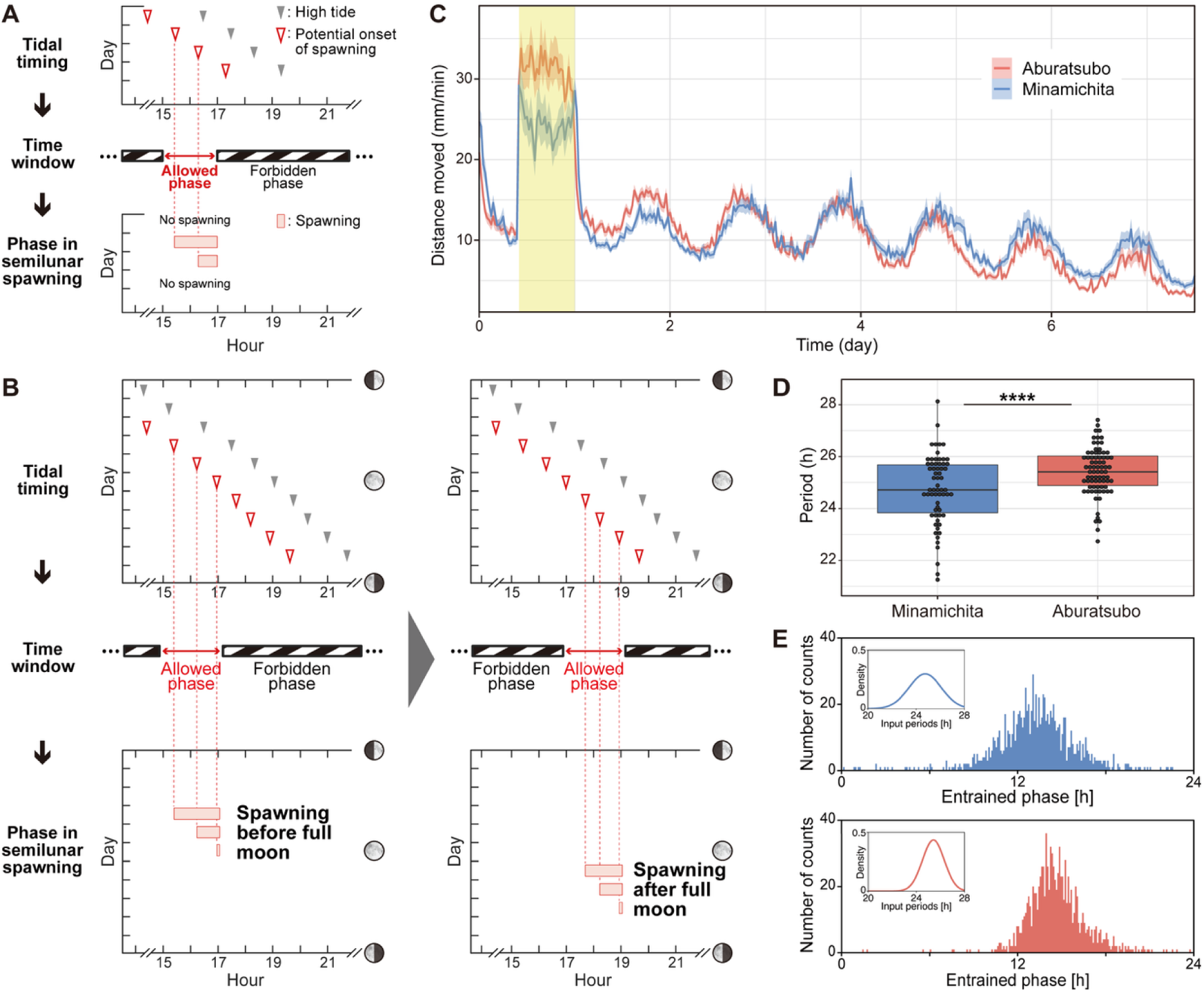
Circadian gating links the spawning time window to the phase of the semilunar spawning rhythm. (A) A schematic diagram of the circadian gating model in grass puffer spawning. Grass puffer spawning occurs a few hours before high tide. In the circadian gating model, however, the circadian clock restricts spawning to a defined daily time window (allowed phase). Spawning occurs only when the potential onset of spawning coincides with this allowed phase. (B) The phase of the spawning time window determines the phase of the semilunar spawning rhythm. Left, Relationship among the potential onset of spawning relative to high tide (top), the spawning time window (15:00-17:00, middle), and the spawning dates (bottom) for Minamichita populations. Right, Replacing the Minamichita spawning time window with that of the Aburatsubo populations (17:00-19:00) shifts the predicted semilunar spawning phase to the latter half of the spring tide. (C) Temporal changes in the average activity levels of larvae in each population (MC: n = 63; AB: n = 77) analyzed in summer 2022. Solid lines and the shaded areas represent the average values and their 95% bootstrap confidence intervals, respectively. The yellow background represents light period. (D) The free-running period of larvae from the two populations calculated using the Lomb-Scargle periodogram (MC: τ = 24.74 ± 1.34 h, n = 63; AB: τ = 25.41 ± 0.91 h, n = 77; Welch’s *t*-test: *t*_(105.53)_ = 3.39, *p* < 0.001). (E) Entrained phase resulting from variation in the endogenous period. Insets show normally distributed endogenous periods (mean ± SD): 24.74 ± 1.34 h (blue, Minamichita) and 25.41 ± 0.91 h (red, Aburatsubo), corresponding to the experimental results shown in Fig. 2D. Simulations of the Poincaré oscillator, an amplitude-phase model^20^ were performed using 1000 endogenous periods sampled from the corresponding normal distributions. The simulated entrained phases were 13.20 ± 2.78 h for Minamichita (blue), and 14.64 ± 2.02 h for Aburatsubo (red). The mean phases differed by 1.44 h between the two populations (Watson–Williams test, *p* < 0.001)

Since the free-running circadian period (τ) is known to influence circadian phase^16^, our model predicts that τ differs between western and eastern populations if variation in the spawning time window underlies geographic differences in semilunar spawning timing. To test this, we next compared the τ of grass puffer from Minamichita and Aburatsubo. In general, locomotor activity rhythms of adult fish lack robustness, but those of larvae are highly robust. Consequently, recording larval behavioral rhythms is well established method in circadian research on fish^17,18^. In order to compare the circadian rhythmic properties, specifically the free-running period, we produced grass puffer larvae from Minamichita and Aburatsubo through artificial fertilization. Hatched larvae were subjected to behavioral assays using a high-throughput measurement system for fish larvae (DanioVision, Noldus). In this assay, larvae were exposed to one light/dark cycle to assess their entrainment to the light and then transferred to constant darkness to evaluate their free-running circadian periods. This experiment confirmed both entrainment to the light/dark cycle and free-running activity under constant dark conditions (Fig. 2C). Analysis using the Lomb-Scargle periodogram revealed that the free-running period of Minamichita (MC) was significantly shorter than that of Aburatsubo (AB) by ∼40 minutes (Fig. 2D) (Welch’s *t*-test: *t*_(105.53)_ = 3.39, *p* < 0.001, n = 63 for MC and n = 77 for AB). These results were consistently replicated in the following year, with the Minamichita population exhibiting a significantly shorter period by ∼70 minutes (SI Appendix, Fig. S1) (Welch’s *t*-test: *t*_(144.95)_ = 9.09, *p* < 0.001, n = 83 for each group). These findings align with the human chronotype research, where a shorter free-running period leads to earlier chronotype (i.e. an earlier time window)^16^.

Although the observed difference in the free-running period between western and eastern populations was 40-70 minutes, the difference in the spawning time window was approximately 2 hours. This discrepancy, in which a relatively small difference in τ results in a larger phase difference, has been reported in previous studies of circadian timing. For example, heterozygous τ mutant hamsters with an approximately two-hour shorter free-running period exhibited activity onsets occurring four or more hours before lights-off^19^. Similarly, a human subject with familial advanced sleep–wake syndrome showed only ∼1 hour shortening of the free-running circadian period but a 3–4 hours advance in sleep phase^16^. A previous theoretical study provides a mechanistic explanation for these phenomena^20^. Using amplitude-phase models, this study demonstrated that the “strong oscillator” property of vertebrate circadian clocks can amplify small variations in τ into larger variations in circadian phase. To determine whether this model could explain our observations, we performed simulations using the τ distributions obtained from behavioral experiments in the Minamichita and Aburatsubo populations (Fig. 2D). The predicted circadian phases by the simulations were 13.20 ± 2.78 h and 14.64 ± 2.02 h for Minamichita and Aburatsubo, respectively (Fig. 2E). Thus, the ∼2 hours difference in spawning time window can be explained by the observed 40-70 minutes difference in τ. Taken together, our field observations, behavioral experiments, and amplitude-phase model simulations indicate that the geographic difference in semilunar spawning phases is likely driven by geographic differences in the circadian phase of the spawning time window.

### Identification of candidate genes by population genomics

Next, to identify the candidate genes responsible for the geographic difference in the spawning time windows, we conducted a population genomics study. Fin samples were collected from a total of 482 individuals across seven populations (three western, three eastern and one outgroup) (Fig. 1A), and genomic DNA was extracted. We pooled the genomic DNA for each population (49–100 individuals per pool) (Fig. 1A). Using paired Illumina NovaSeq S4 reads, we performed both pooled and individual DNA whole-genome sequencing with an average depth of 28× coverage. This resulted in 6.3 M SNPs being retained after stringent filtration. A principal component analysis (PCA) plot of individuals from the seven populations showed distinct patterns among the western, eastern populations and the outgroup, suggesting genetic differentiation between the western and eastern populations (Fig. 3A).

**Fig. 3.**
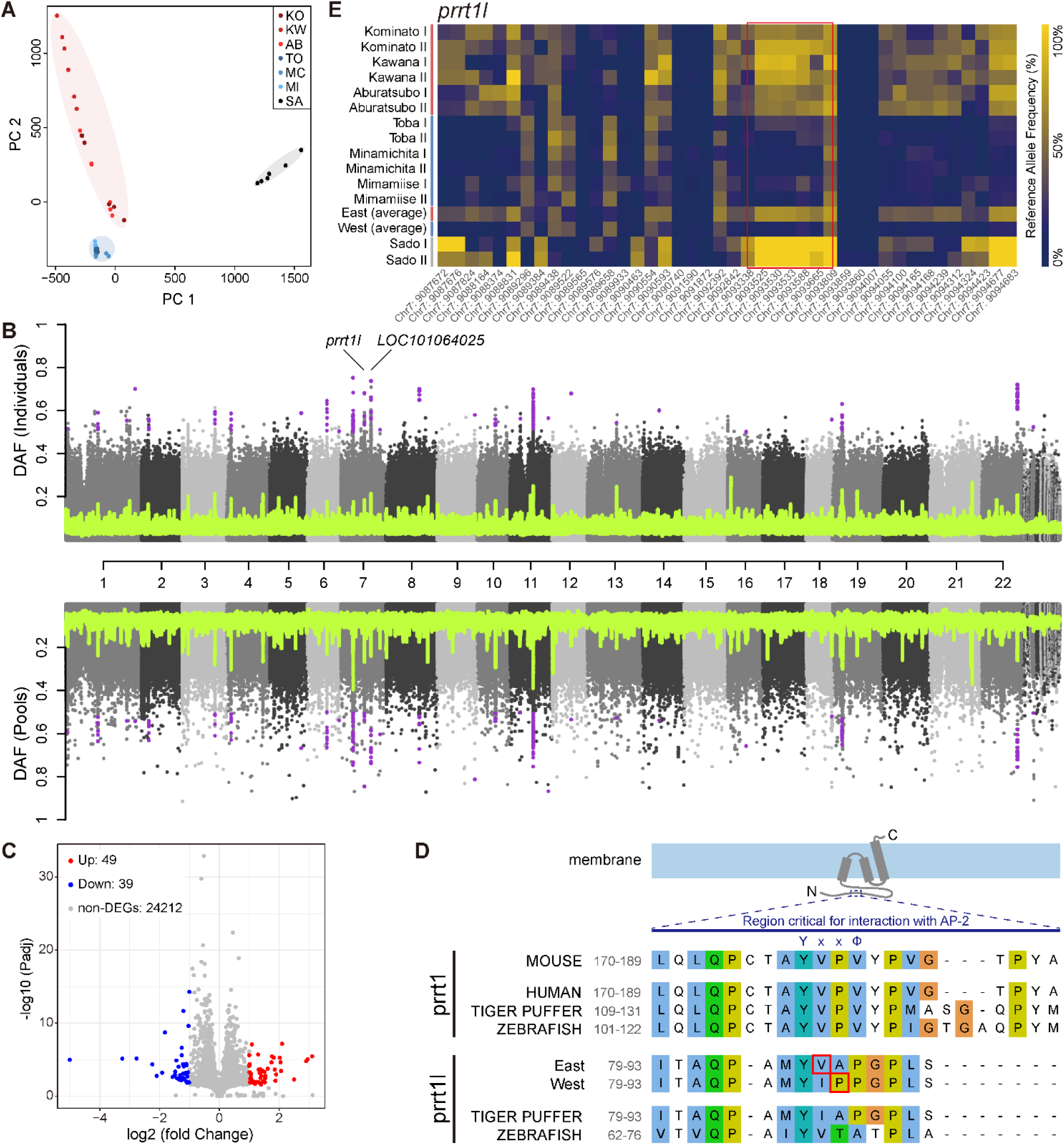
Population genomics identified *prrt1l* as a candidate gene. (A) A principal component analysis (PCA) plot of genome sequences in the seven populations. The blue, red and grey ellipses represent the western (Minamichita, Toba, and Minamiise), eastern (Aburatsubo, Kominato, and Kawana) and outgroup (Sado) populations, respectively. (B) Delta allele frequency (DAF) contrast between western and eastern populations. The gray dots show single SNPs, the green lines a 100 SNP rolling average. (C) Identification of 88 DEGs between the western (Minamichita and Toba) and the eastern (Aburatsubo and Kominato) populations by RNA-seq of the hypothalamus and pituitary. (D) Schematic representation of the prrt1/prrt1l protein and a multiple sequence alignment of its interaction region with AP-2. The cytoplasmic N-terminal domain contains the region that is critical for interaction with the AP-2–interacting segment reported for mouse PRRT1^28^. Yxxɸ (ɸ: a hydrophobic residue) is a canonical AP-2 sorting signal. Sequences from mouse, human, zebrafish, tiger puffer and western and eastern populations of grass puffer are compared. Residue numbers on the left corresponds to the position in the full-length protein. Amino acid substitutions in the western and eastern populations are highlighted with red boxes. The displayed region was prepared in Jalview using the Clustal color scheme. (E) A zoomed-in heatmap of the genomic region of *prrt1l* (Chr7: 9087672–9094683 bp), indicated by the red arrowhead. Regions with strong differentiation are highlighted with a red box.

We then performed a genome-wide compilation of independent regions of differentiation (Fig. 3B). We estimated differentiation between the western and eastern populations and identified highly differentiated regions. In these regions, 24 genes were located, and we considered them as potential candidate genes (SI Appendix, Table S1). Comparison of the genomic DNA sequences for these 24 genes between the western and eastern populations revealed missense variants that caused amino acid substitutions in two genes including proline rich transmembrane protein 1-like (*prrt1l*) gene and high-affinity choline transporter 1-like (*LOC101064025*) (Fig. 3B).

Lunar-regulated beach-spawning behaviors are considered to be regulated by the hypothalamic-pituitary regions^10^. To further identify genes with differential expression levels among the 24 candidate genes, we performed RNA sequencing (RNA-seq) analysis using the hypothalamic-pituitary brain regions collected in the field during the spawning time from the western (Minamichita and Toba) and eastern (Aburatsubo and Kominato) populations. For each population, two individuals were pooled per biological replicate (eight fish in total, n = 4 each), and RNA-seq was performed using the DNB platform, generating an average of 5.59 Gb of sequence data per sample. Using DESeq2 algorithms, we identified 88 differentially expressed genes (DEGs) with greater than 2-fold changes (Fig. 3C and SI Appendix, Table S2). Interestingly, Gene Ontology (GO) classification of the 88 DEGs revealed *per3* as a gene in the “rhythmic process” category, which may reflect the difference in rhythmic phenotype between the two populations (SI Appendix, Figs. S2 and S3). Its expression was higher in the western populations (adjusted *P*-value < 0.001, SI Appendix, Fig. S3). However, importantly, none of the 24 candidate genes was differentially expressed between the two populations.

Based on these results, we prioritized prrt1l and LOC101064025, which exhibited amino acid substitutions between the western and eastern populations. Upon checking our RNA-seq results, we found that *prrt1l* was expressed, whereas *LOC101064025* was not, in the hypothalamic-pituitary regions (SI Appendix, Fig. S4). We therefore focused on *prrtl1* gene, which exhibits missense mutations between the western and eastern populations yielding an isoleucine for valine substitution in the eastern population and an alanine for proline substitution in the western population compared with the closely related tiger puffer (*Takifugu rubripes*) (Fig. 3D). Inspection of allele frequencies for SNPs in the potential candidate genes revealed significant differences between the western and eastern populations at *prrt1l* locus (Fig. 3E).

### Expression of *prrt1l* and circadian clock genes in the suprachiasmatic nucleus

Prrt1/prrt1l belong to the CD225 (dispanin) superfamily, whose members regulate vesicular membrane fusion^21,22^. It is also a component of the native AMPA-type glutamate receptor (AMPAR) complex, modulating its function^22^. A recent study employing machine learning-based feature selection methods (i.e., Lasso logistic regression and Support Vector Machine-Recursive Feature Elimination) reported significant Pearson correlations between the expression levels of *PRRT1* and circadian clock genes *CLOCK*, *PER1*, *PER3*, and *CRY1*, the strongest of which was with *PER3*^23^. It is interesting to note that *per3* showed differential expression in our RNA-seq analysis between western and eastern populations (SI Appendix, Fig. S3). In addition, *per1* and *cry1l* exhibited smaller yet significant changes (adjusted *P* < 0.05; log2FC = −0.38 and 0.65, respectively), though they did not pass our DEG threshold (|log2FC| ≥ 1) (SI Appendix, Fig. S3).

To examine the localization of the grass puffer *prrt1l* and clock gene expressions, we reanalyzed our spatial transcriptome dataset^10^. The brain region containing the hypothalamus and pituitary was clustered into eight clusters, and the suprachiasmatic nucleus (SCN), the circadian pacemaker, was mapped to cluster 3 (Fig. 4A and B). Expression of *prrt1l* and clock genes was detected within the SCN (Fig. 4C). These observations are consistent with a model in which *prrt1l* is correlated with the circadian clock genes^23^. We therefore hypothesized that knocking out *prrt1l* might affect free-running rhythms.

**Fig. 4.**
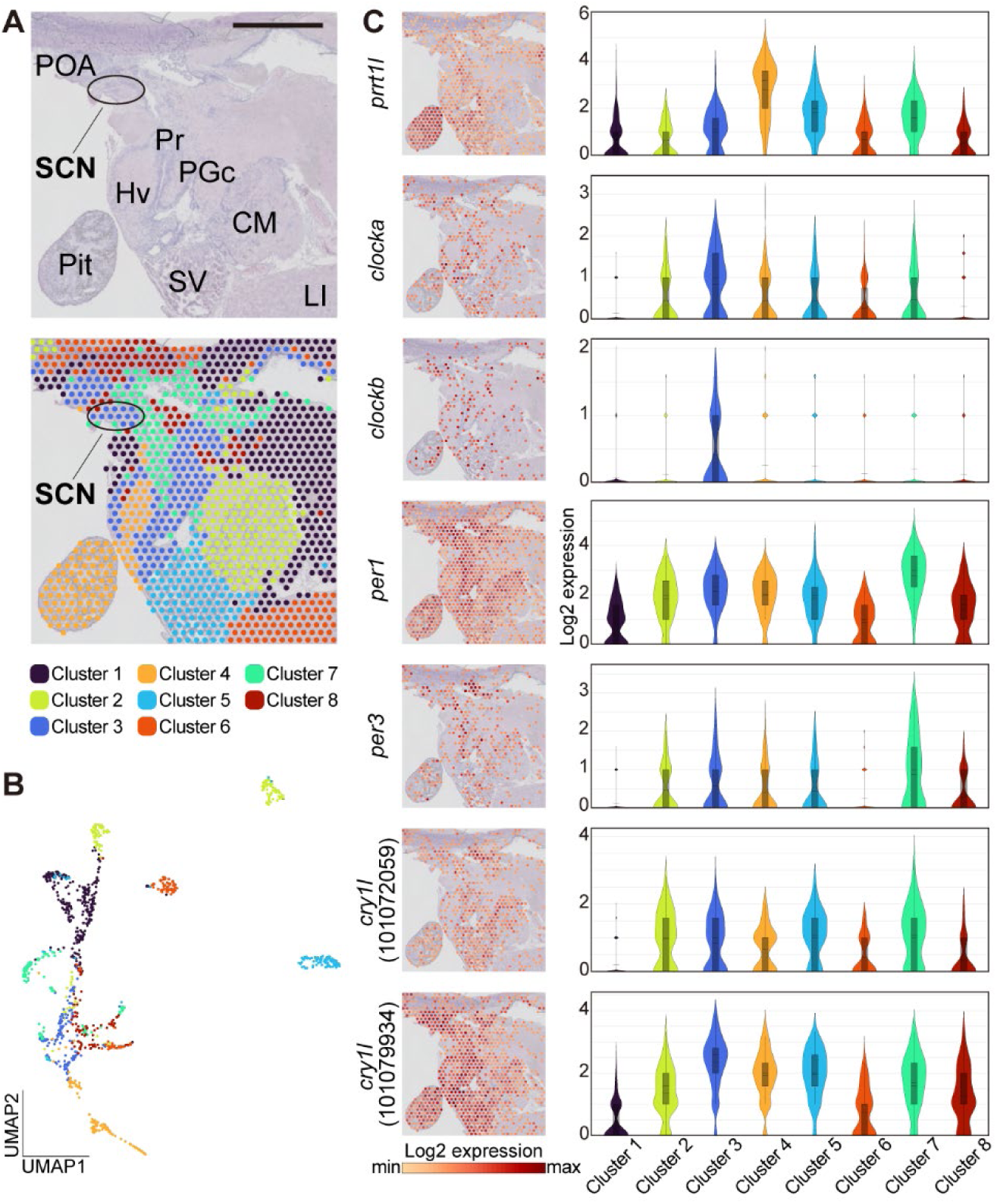
Localization of *prrt1l* and circadian clock gene expression, revealed by spatial transcriptomics of the grass puffer hypothalamus and pituitary. (A) The brain regions used in this analysis and the spatial distribution of the eight clusters. The suprachiasmatic nucleus (SCN) belongs to cluster 3. Scale bar: 1 mm. (B) UMAP showing the eight clusters. (C) Spatial expression patterns and violin plots showing the expression levels of *prrt1l* and the circadian clock genes that are reported to be associated with PRRT1 in Zhang et al. (2025)^23^. Abbreviations: SCN: suprachiasmatic nucleus; POA: anterior parvocellular nucleus of preoptic nucleus; Pr: periventricular region of diencephalon; PGc: commissural preglomerular nucleus; Hv: ventral zone of periventricular hypothalamus; Pit: pituitary; CM: corpus mammillare; SV: saccus vasulosus; LI: lobus inferior.

### Shorter free-running period in *prrt1l* F0 knockout larvae

To test the functional significance of *prrt1l* in free-running activity rhythms, we generated *prrt1l* knockout grass puffer individuals for behavioral analysis. With this aim, we employed a triple CRISPR F0 knockout approach^24,25^. This method uses three guide RNAs (gRNA) per gene to achieve highly efficient knockout without the need for breeding, thus enabling reverse genetic analysis even for wild animals such as grass puffer, which are difficult to raise to adulthood in a laboratory setting. We designed three gRNAs that target the first, second and third exons of *prrt1l.* We then injected these gRNAs, along with Cas9 protein, into eggs that were obtained by artificial fertilization at Aburatsubo (Fig. 5A).

**Fig. 5.**
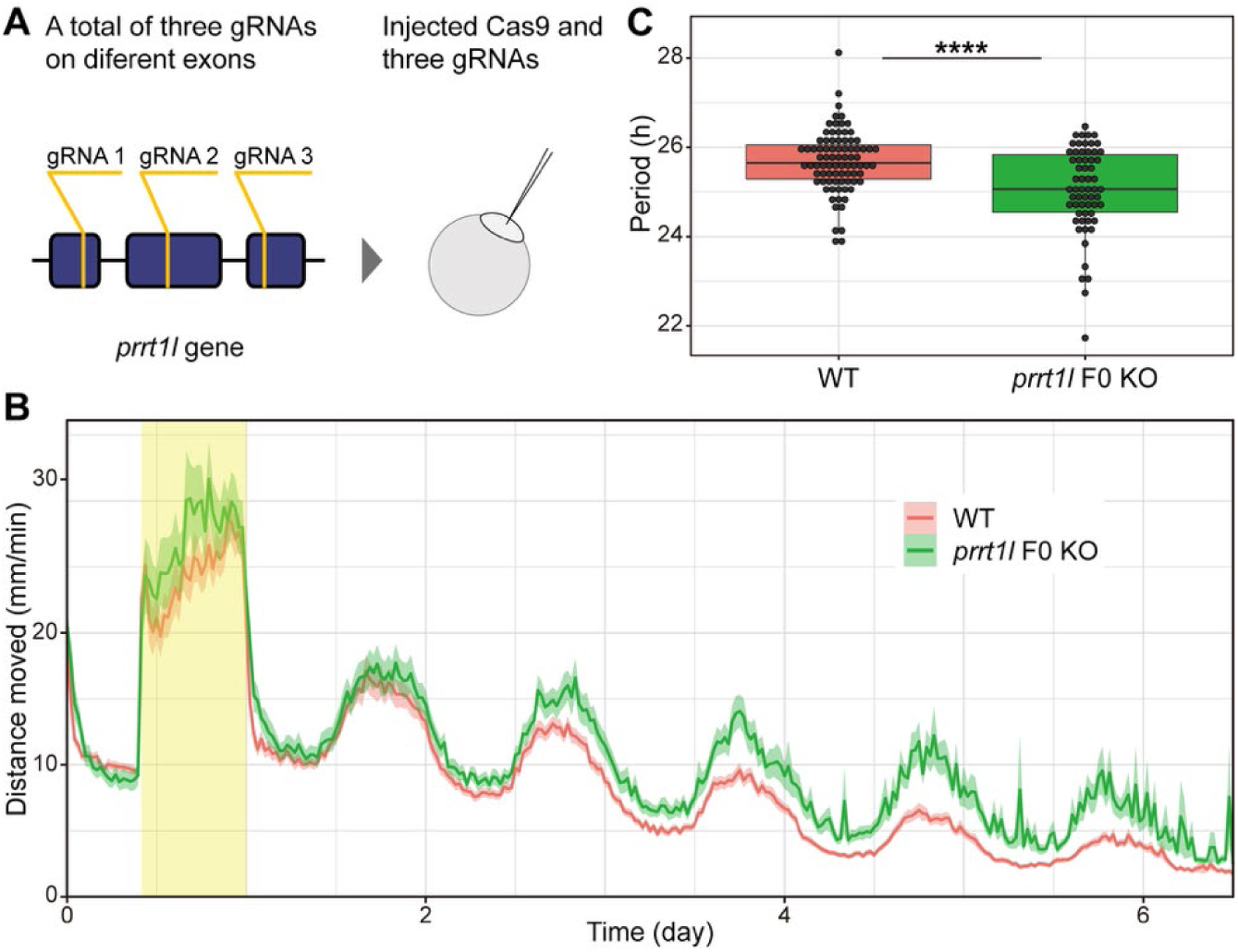
*prrt1l* F0 knockout shortened the free-running rhythm in grass puffer larvae. (A) A schematic diagram of triple CRISPR in puffer larvae. (B) Temporal changes in the average activity levels of larvae in wild-type (WT) and *prrt1l* F0 knockout (KO) larvae. The solid lines and shaded areas represent the average values and their 95% bootstrap confidence intervals, respectively. The yellow background represents a light period. (C) Free-running periods of WT and *prrt1l* F0 KO larvae, as calculated using the Lomb-Scargle periodogram (WT: τ = 25.68 ± 0.69 h, n = 83; *prrt1l* F0 KO: τ = 25.24 ± 0.96 h, n = 63; Welch’s *t*-test: *t*_(176.42)_ = 3.42, *p* < 0.001).

Using the larvae that hatched from these injected eggs, we conducted a behavioral analysis similar to that used to compare the western and eastern populations in Fig. 1. Following the experiment, we genotyped the larvae and only used data from biallelic mutant individuals. The behavioral analysis revealed that both WT and *prrt1l* F0 KO individuals exhibited entrainment to the light/dark cycle and free-running activity under constant dark conditions (Fig. 5B). Notably, the free-running period of *prrt1l* F0 KO individuals was significantly shorter than that of WT by ∼30 minutes (Fig. 5C) (Welch’s *t*-test: *t*_(176.42)_ = 3.42, *p* < 0.001; n = 83 for WT and n = 63 for *prrt1l* F0 KO).

## Discussion

The genetic basis underlying the lunar rhythms remains unclear. The timing of the grass puffer spawning is synchronized with high tide during the spring tide. In the present study, we demonstrated that western populations spawn during the first half of the spring tide, whereas eastern populations spawn during the second half. Our analysis also showed that puffer spawning was restricted to narrow daily time window, with western populations exhibiting an earlier circadian phase than eastern populations (Fig. 1). Since the free-running circadian period (τ) influences circadian phase^16^, we compared the τ between grass puffer populations from Minamichita and Aburatsubo. We then found that the Minamichita population had a significantly shorter free-running period than the Aburatsubo population. Both the circadian gating model and mathematical simulations suggest that the free-running period (τ), the phase of the spawning time window, and the phase of semilunar rhythm are mechanistically linked (Fig. 2). Thus, the natural variation in lunar-synchronized spawning rhythms between western and eastern grass puffer populations provides an excellent model for understanding the genetic basis of lunar rhythms.

The population structure of coastal fish around the main islands of Japan is influenced by the Kuroshio and Tsushima currents (Fig. 1A), resulting in Pacific Ocean and Sea of Japan lineages of fish species^26^. A PCA plot of individuals from the puffer populations revealed distinct patterns between western and eastern populations (Fig. 3A). The geographic boundary of this variation lies around the Izu Peninsula (Fig. 1A). Interestingly, the genotypes of the surfperch *Ditrema jordani* are also divided between western and eastern Japan by the Izu Peninsula. It has been hypothesized that the unique reproductive characteristics of *D. jordani,* specifically viviparity and the birth of fully developed juveniles, limit dispersal and result in a strong local population structure^27^. In the grass puffer, high specificity in spawning sites and homing behavior likely contribute to local population structure. Furthermore, differences in spawning timing may have reinforced genetic differentiation, potentially acting as a partial reproductive barrier. Understanding the ecological and behavioral factors affecting population structure is crucial for grasping how diversity is generated^26^; thus, the grass puffer can also serve as a valuable model in phylogeography.

In the present study, we employed unbiased, discovery-driven approaches such as a population genomics approach and a transcriptome analysis, to identify the responsible genes. The reproduction of the marine midge *Clunio marinus* is also timed by circadian and circalunar clocks, and it has been suggested that calcium/calmodulin-dependent kinase II.1 (CaMKII.1) is involved in timing adaptation^9^. Unlike the marine midge study, we did not identify any circadian clock-related genes, but we did identify *prrt1l* as a potential modulator of the circadian period in the grass puffer through population genomics analysis and F0 KO larvae analysis (Figs. 3 and 5). We also observed the expression of *prrt1l* and circadian clock genes in the grass puffer SCN by spatial transcriptomics (Fig. 4).

There are several possible mechanisms by which prrt1l affects circadian periods. Of particular interest is the fact that a machine learning approach suggested the correlation between *PRRT1* and circadian clock genes, especially *PER3*, *CRY1*, and *PER1*^23^. Consistent with this, our RNA-seq analysis revealed differential expression levels of the *per3, cry1l,* and *per1* genes between the western and eastern populations (SI Appendix, Fig. S3). Therefore, *prrt1l* may regulate the circadian period by altering the expression of circadian clock genes.

A recent study reported that the PRRT1 protein in mice interacts with the µ2 subunit of the adaptor protein complex AP-2, which regulates endocytic trafficking^28^. Notably, the amino acid substitutions we identified in *prrt1l* are located within a region homologous to the reported AP-2-interacting site (Fig. 3D), suggesting that the geographic variation in *prrt1l* may alter endocytic trafficking. Growing evidence highlights interactions between circadian rhythms and membrane trafficking pathways. For example, *Rab3a* (a regulator of synaptic vesicle trafficking) mutant mice called ‘earlybird’ exhibit a shorter free-running period and an advanced chronotype^29^. In humans, two independent GWAS for chronotype identified ERC2, which belongs to the Rab3-interacting molecule-binding protein family, involved in neurotransmitter release^30–32^. Similarly, CAVIN3, a factor associated with caveolae (microdomains of the plasma membrane), regulates the circadian period^33^. Collectively, these observations suggest potential crosstalk between circadian rhythms and membrane trafficking.

It is also noteworthy that prrt1 is a component of the native AMPA-regulated glutamate receptor complex, thereby modulating glutamate signaling^22^. AMPA receptors are involved in the photoentrainment of the circadian clock mediated by the neurotransmitter glutamate^34^. Interestingly, a recent genome-wide association study on temporal niches in cichlids identified *nsg2* as a candidate gene near a top-ranked locus for crepuscularity; this protein binds to AMPA receptors and regulates their surface expression^35,36^. In the present study, however, we did not observe significant behavioral changes in response to the light/dark cycle in F0 KO and wild-type larvae (Fig. 5). Due to the technical limitations, we were unable to evaluate the effects of twilight on the behavioral analyses of puffer larvae. The behavior of animals under artificial light dark conditions can differ significantly from that observed under natural conditions^37,38^. Therefore, examining the behavior of F0 KO larvae under natural conditions in the future would also be of great interest.

Although prrt1 is known to be involved in glutamate signaling and suggested to correlate with circadian clock genes^22,23^, current knowledge regarding the function of prrt1 remains very limited. Phylogenetic analysis revealed that prrt1 and prrt1l form close but distinct clades within the CD225 protein family (SI Appendix, Fig. S5). Therefore, it is also possible that prrt1l may regulate the circadian rhythms through a mechanism distinct from the functions previously identified in prrt1.

Although the precise molecular mechanism by which prrt1l regulates circadian rhythms remains unclear, unbiased genetic approaches frequently provide novel insights. Future studies into prrt1l are therefore expected to uncover previously unknown regulatory mechanisms governing both circadian and lunar rhythms.

### Limitation of the study

The grass puffer is a wild animal, and it remains technically impossible to raise KO animals until they reach adulthood. Furthermore, adult grass puffers do not exhibit reproductive behavior and spawning in captivity as they do in the wild. This is consistent with previous findings that female tiger puffers fail to undergo normal oocyte maturation in captivity^39,40^.

## Materials and methods

### Animals

For the population genomics analysis, fins were collected from adult grass puffers across seven different populations (Fig. 1A). For the behavioral assay of larvae, puffer embryos were obtained by artificial fertilization (as detailed in the following section) at Minamichita and Aburatsubo during the spring tides from mid-May to mid-July. To collect samples for the RNA-seq experiment, adult male puffers were anesthetized with 0.05% ethyl 3-aminobenzoate methanesulfonate (MS-222, Sigma-Aldrich) and brain tissues containing the hypothalamus and pituitary were collected. All animal experiments were approved by the Animal Experiment Committee of Nagoya University and carried out in accordance with ARRIVE guidelines.

### Artificial fertilization using the wild grass puffer

Mature male and female grass puffers were caught during the spring tides. In the natural environment, the male-to-female ratio in the spawning group is much higher for males, typically 4–40 times higher. To ensure genetic diversity in the fertilized eggs, at least three females and many more males were collected. The sperm and eggs were gently pushed out into a bowl, mixed together, and left to stand for one minute. Subsequently, seawater was added to the bowl and left to stand for another minute. The eggs were then washed in seawater until the supernatant became clear. Fertilized eggs were kept in artificial seawater (SEALIFE, Marinetech) until hatching under long-day and warm conditions (14 hours of light and 10 hours of darkness at 22℃) corresponding to natural conditions during the breeding season.

### Data collection on spawning timing

Field observations were conducted at a known spawning site in Minamichita. In contrast, since no spawning sites had been previously reported in Toba, preliminary surveys were performed and identified a spawning site. As spawning was visible from the shore at both sites, observations were conducted from the shore during spring tides to record the times of the first and last observed spawning events. Data on spawning times at Aburatsubo and Kominato were obtained from previous studies to facilitate comparison^11,14^. Tide tables for each spawning site were obtained from the tide736.net (https://tide736.net/), a website offering tide data based on official calculations by the Japan Coast Guard.

### Population genomics

Tail fins of grass puffer were sampled from 49–100 individuals per population and stored at −80 °C. DNA extraction from the samples was conducted using DNeasy Blood and Tissue Kits (Qiagen). Genomic DNA of 49–100 individuals per population were pooled at equal concentrations for pooled DNA whole-genome sequencing (pool-seq), while 14 individuals (seven females and seven males) from Minamichita and six individuals from each other location was picked out of individual sequencing.

### Whole-genome sequencing, mapping and genotyping

Illumina short-read sequencing of pooled and individual genomic DNA was carried out using the standard configuration of paired 150 bp reads from a random shotgun library generated by enzymatic fragmentation followed by size selection, performed at NGI Uppsala—SNP&SEQ Technology Platform. Data were generated using one lane of Illumina NovaSeq S4, comprising 2.5 × 10^9^ read pairs: For pools the range per sample was 2.3 – 6.3 × 10^7^; for individuals it was 1.7 – 8.3 × 10^7^. The reads are available at NCBI’s Short Read Archive (BioProject: PRJNA1377317). Reads were mapped to fTakRub1.2 (GCF_901000725.2) using bwa mem v 0.7.17-r1188^41^, and PCR duplicates were marked using “MarkDuplicates” from Picard Tools (http://broadinstitute.github.io/picard/). The resulting bam files were then processed using GATK v4.1.1.0^42^, with the following workflow: First, we used “HaplotypeCaller” to generate per-sample gvcf files. These were merged using “CombineGVCFs” and genotyped using “GenotypeGVCFs.” The raw SNP genotypes were then passed through the “VariantFiltration” module, with the following arguments: --filter-expression "QD < 8.0 || FS > 50.0 || MQ < 30.0 || MQRankSum < −10.0 || ReadPosRankSum < −6.0" --missing-values-evaluate-as-failing using --filter-expression "DP < 100.0 || DP > 1500". After this procedure, >97 % of reads were mapped, with a 2.4 % mismatch rate.

### Delta allele frequency contrasts and heatmaps

Frequencies were calculated either from genotypes (individuals) or the REF (the allele in the reference genome) and ALT (any other allele found at that locus) read counts (pools) at each SNP position. These frequencies were subsequently used in heatmaps and Delta allele frequency (DAF) contrasts. For both individual and pool contrasts, the samples were grouped according to the division in Fig. 1, with samples from Aburatsubo, Kawana and Kominato constituting the eastern group, and those from Minamiise, Minamichita and Toba constituting the western group. All contrasts were based on the absolute (i.e non-signed) DAF of the groups involved. We calculated DAF both on a per-SNP basis, and as a rolling average across 100 adjacent SNPs, and defined SNPs that had DAF > 0.5 in both contrasts as concordant high-differentiation SNPs.

### RNA-seq analysis

RNA-seq analysis was performed on the hypothalamus and pituitary of fish from two western populations (Minamichita and Toba) and two eastern populations (Aburatsubo and Kominato). Adult male puffers were collected at their spawning site on the days of spawning. Total RNA was extracted from the hypothalamus and pituitary using a RNeasy Lipid Tissue Mini Kit (Qiagen) combined with a Multi-beads shocker (Yasui Kikai). For each population, four biological replicates were prepared by pooling equal amounts of RNA from two individuals. mRNA was purified using poly(dT) oligo-attached magnetic beads, followed by fragmentation and subsequent cDNA synthesis. The synthesized cDNA was then used to prepare PE100 strand-specific sequencing libraries, which were subsequently sequenced using a DNBseq platform to generate 50–60 million reads for each library (BGI). Clean reads were first aligned to the fTakRub1.2 reference genome using HISAT2 v2.0.4^43^. Transcripts were then assembled and compared to the reference to identify novel transcripts, which were integrated in order to construct an updated reference. The reads were then re-aligned to this updated reference with Bowtie2 v2.2.5^44^, and gene-level abundances were estimated with RSEM v1.2.12^45^. Differential expression was tested using DESeq2 with thresholds of an adjusted *p*-value ≤ 0.05 and |log2 fold change| ≥ 1 to define differentially expressed genes (DEGs)^46^. DEGs were classified by Gene Ontology (GO) annotation.

### Spatial transcriptomics

Spatial transcriptomics data of the grass puffer brain were obtained from our previous study^10^. To focus on the suprachiasmatic nucleus (SCN), the hypothalamic region was sub-selected and reanalyzed using the Reanalyze workflow in Loupe Browser v8.1.2 (10x Genomics), totaling 1134 spatial barcodes. Clustering was performed based on 10 principal components, and UMAP was computed with a minimum distance of 0.2 and 20 neighbors.

### Behavioral assay of larvae

Hatched larvae (8–9 days post-fertilization (dpf)) were placed in each well of a 96-well plate (4titude) containing 800 µl of artificial seawater (SEALIFE, Marinetech). The plate was placed in a DanioVision (Noldus) at 22 ℃. A CCD camera and an infrared light source were used for recording. For the first 1-2 days, the larvae were kept under long-day condition (14 hours of light and 10 hours of darkness) consistent with their hatching condition to observe their entrainment to the light/dark cycle. They were then transferred to constant darkness for a week. Following the experiment, the larvae were anesthetized with 0.05% MS-222 for genotyping. Tracking data were processed using EthoVision XT14 (Noldus), and behavioral data for each individual were calculated at one-minute intervals. The raw data were processed using R and the R package Rethomics^47^. The free-running period was calculated using the Lomb-Scargle periodogram (period range = 18–30) on the behavioral data between 2.5‒7 days under constant darkness.

### Generation of triple CRISPR F0 KO larvae

The F0 knockout protocol for zebrafish was used with some modifications^25^. Candidate gRNA sequences were designed using CHOPCHOP, with fTakRub1.2 as a reference^48^. To confirm sequence conservation, these candidates were cross-referenced with RNA-seq data from both western and eastern populations. Consequently, three gRNAs targeting three different exons were selected (SI Appendix, Table S3). The synthetic gRNAs (composed of two components: custom crRNA and tracrRNA) were purchased from Integrated DNA Technologies (IDT). Custom crRNAs were designed based on the gRNA sequences via the IDT website (SI Appendix, Table S3). The crRNA was annealed with an equal amount of tracrRNA and diluted to 30.5 μM in Duplex buffer (IDT). The Cas9 protein (IDT) was diluted to 30.5 μM with Duplex buffer and mixed with an equal amount of the annealed gRNA to generate a 15.75 μM RNP solution. The three RNP solutions targeting the three different exons were then pooled in equal amounts prior to injections. For F0 KO of grass puffers, embryos were obtained by artificial fertilization at Aburatsubo. Following the experiment, larvae were anesthetized and genotyped. In the genotyping process, a sequence of ∼6 kb, including the whole *prrt1l* gene region, was amplified. Although even a single-base indel could potentially result in gene knockout, we strictly selected only biallelic mutant individuals that did not display the wild-type (WT) amplification pattern for subsequent analyses.

### Protein sequence alignment

The full-length prrt1 and prrt1l protein sequences of mouse, human, zebrafish and tiger puffer were obtained from UniProt and RefSeq, and accession numbers were as follows: O35449 (Mouse PRRT1), Q99946 (Human prrt1), XP_029700676.1 (Tiger puffer prrt1), XP_688192.3 (Zebrafish prrt1), XP_003965976.1 (Tiger puffer prrt1l), NP_001373279.1 (Zebrafish prrt1l). The alignments were generated using MAFFT version 7 with the default parameters and were visualized using Jalview with the Clustal color scheme^49,50^. Only regions homologous to the AP-2–interacting segment are shown in Fig. 3D.

### Phylogenetic analysis

The phylogenetic tree of the tiger puffer prrt1l and its relatives was generated using SHOOT.bio with metazoa database^51^. The tree data were retrieved using R to reduce the number of species and visualized using Interactive Tree of Life^52^.

### Simulations of the amplitude-phase model

Figure 2E was generated using simulations of the Poincaré oscillator, an amplitude-phase model described previously^20^. The model with radius ***r*** and phase ***φ*** is given by

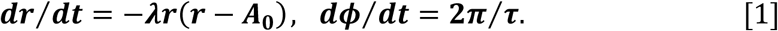

In the model, ***A*_0_** and ***τ*** denote the oscillator amplitude and intrinsic period, respectively, and ***λ*** determines the relaxation rate toward the stable limit cycle (***r*** = ***A*_0_**). Periodic forcing was applied by adding ***F* sin**(**2*πt***⁄***T***) to the *x*-coordinate of the oscillator. Simulations were performed with ***λ*** = **5**. **0**, ***A*_0_** = **1**. **0**, *T* = 24, and *F* = 0.05. The phase after 50 days of forcing is shown, where phase 0 corresponds to the state (*x*,*y*) = (1,0). Numerical simulations were performed using Mathematica (Version 14.3, 2024, Wolfram Research, Inc., Champaign, IL).

### Statistical analysis

Two independent experiments were conducted in the behavioral assays. To compare the free-running periods, a two-tailed Welch’s *t*-test was used to calculate the *p-*values. For circular data, differences in mean phase were assessed using the Watson–Williams test.

## Data availability

The RNA-seq data generated in this study have been deposited in the NCBI’s Gene Expression Omnibus (GEO) (accession numbers GSE313434). The genomic DNA read data are available at NCBI’s Short Read Archive (BioProject: PRJNA1377317).

## Acknowledgements

We thank Drs. Naoyuki Yamamoto and Taeko Nishiwaki-Ohkawa for helpful discussion. This work was supported by the JSPS KAKENHI Grant-in-Aid for Scientific Research (S) 24H00058 (T.Yo.), Grant-in-Aid for Transformative Research Areas (A), Chronoproteinology 24H02303 (T.Yo.), a JSPS

Research Fellowship for Young Scientist program 24KJ1243 (Y.K.). WPI-ITbM is supported by the World Premier International Research Center Initiative (WPI), MEXT, Japan.

## Author contributions

T.Yo. conceived the study. Y.K., D.K., M.P., J.C., L.A., and T.Yo. designed the research. Y.K., D.K., M.P., J.C., L.R., T.Ya., T.N., K.O., M.M., R.E., G.K., L.A., and T.Yo. conducted the experiments and analyzed the data. H.A., A.S. and Y.H. provided new material and methods. Y.K., M.P., G.K., L.A. and T.Yo. wrote the manuscript. All authors discussed the results and commented on the manuscript. T.Yo. supervised the study.

## Declaration of interests

The authors declare no competing interests.

## Supplementary Figures

**Fig. S1.**
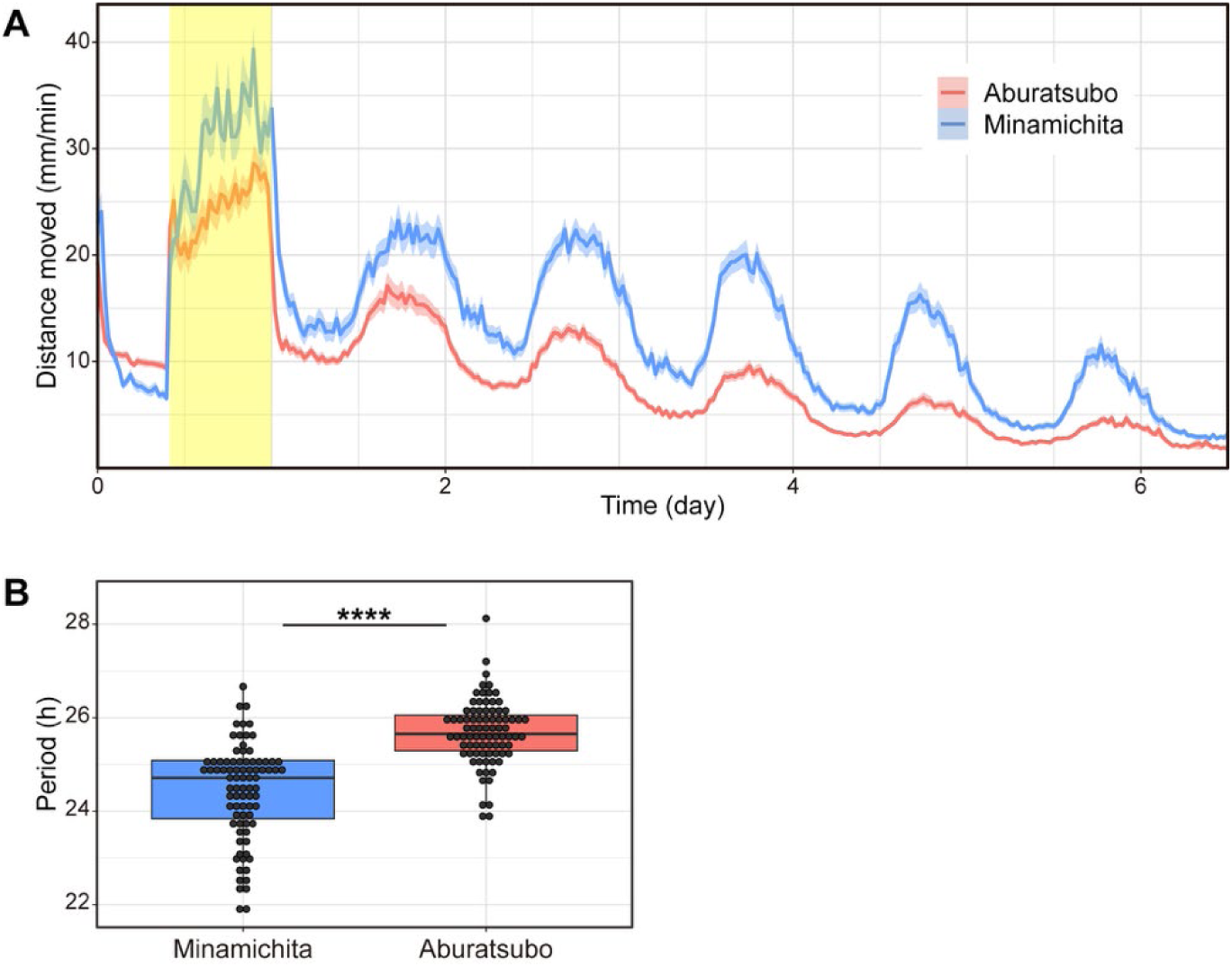
Shorter free-running period in Minamichita than Aburatsubo. (A) Temporal changes in the average activity levels of larvae in each population analyzed in summer 2025. The solid lines represent the average values, and the shaded areas represent the 95% bootstrap confidence interval. The yellow background represents the light period. (B) The free-running period of larvae from two populations calculated using the Lomb-Scargle periodogram (Aburatsubo: τ = 25.68 ± 0.69 h, n = 83; Minamichita: τ = 24.43 ± 1.02 h, n = 83; Welch’s *t*-test: *t*_(144.95)_ = 9.09, p < 0.001).

**Fig. S2.**
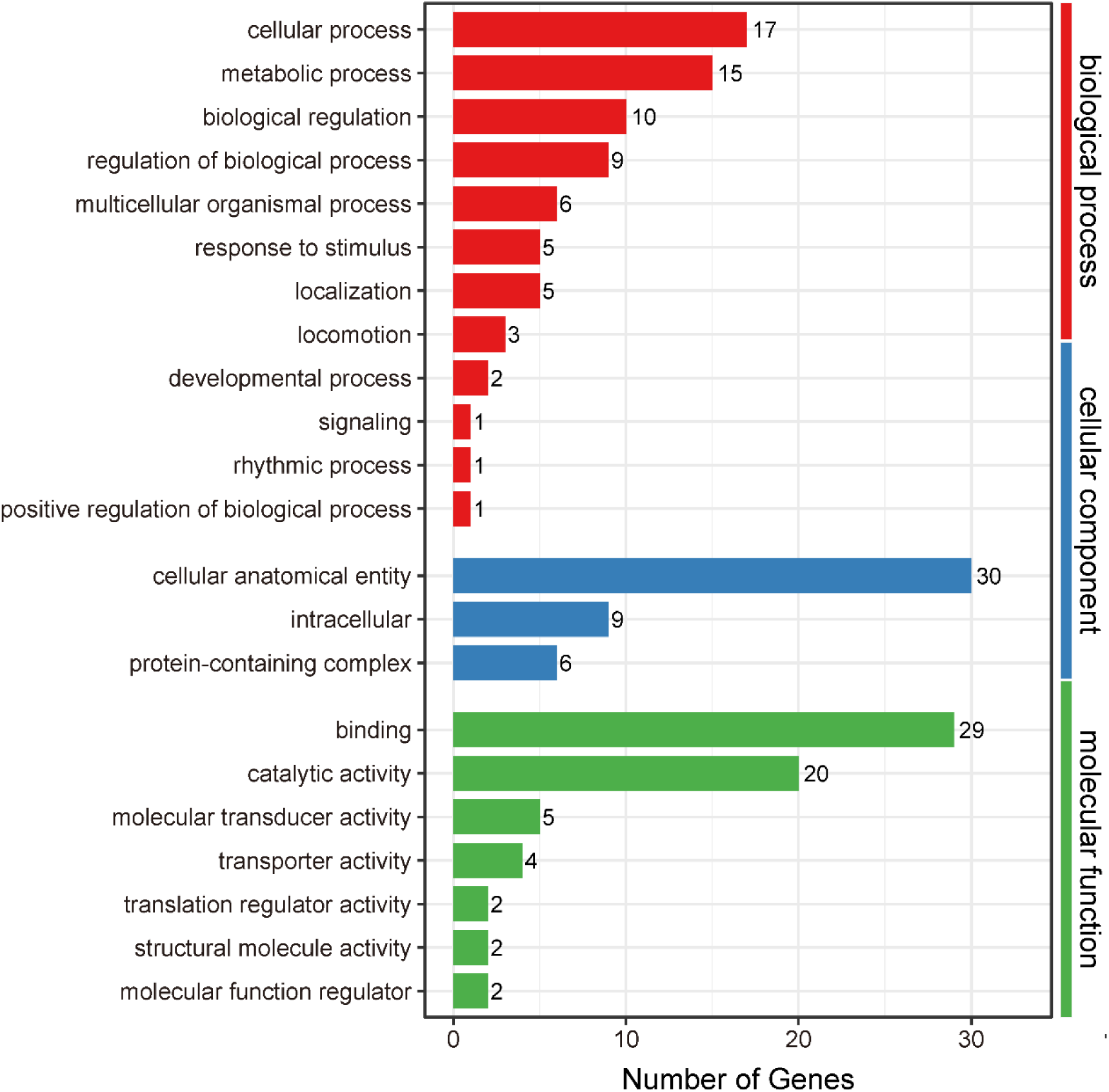
Gene ontology classification of the 88 DEGs. X axis represents number of DEGs. Y axis represents GO term.

**Fig. S3.**
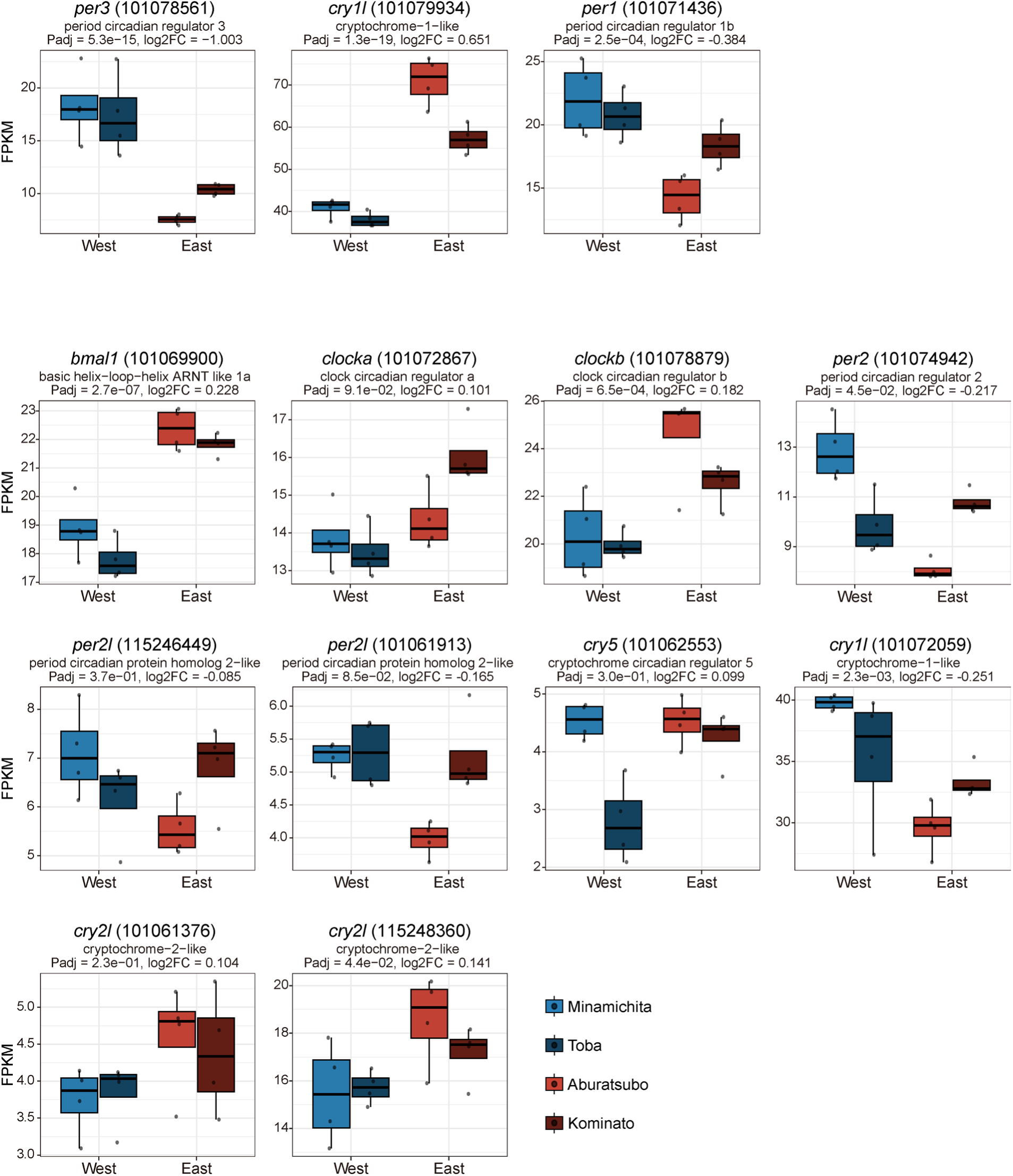
Expression levels of all core clock genes in RNA-seq data from western (Minamichita and Toba) and eastern (Aburatsubo and Kominato) populations. Clock genes exhibiting different expression patterns between populations are shown in the upper panel and other clock genes are shown in the lower panel. Boxplots show the distribution of expression levels in each population, with black dots representing individual values.

**Fig. S4.**
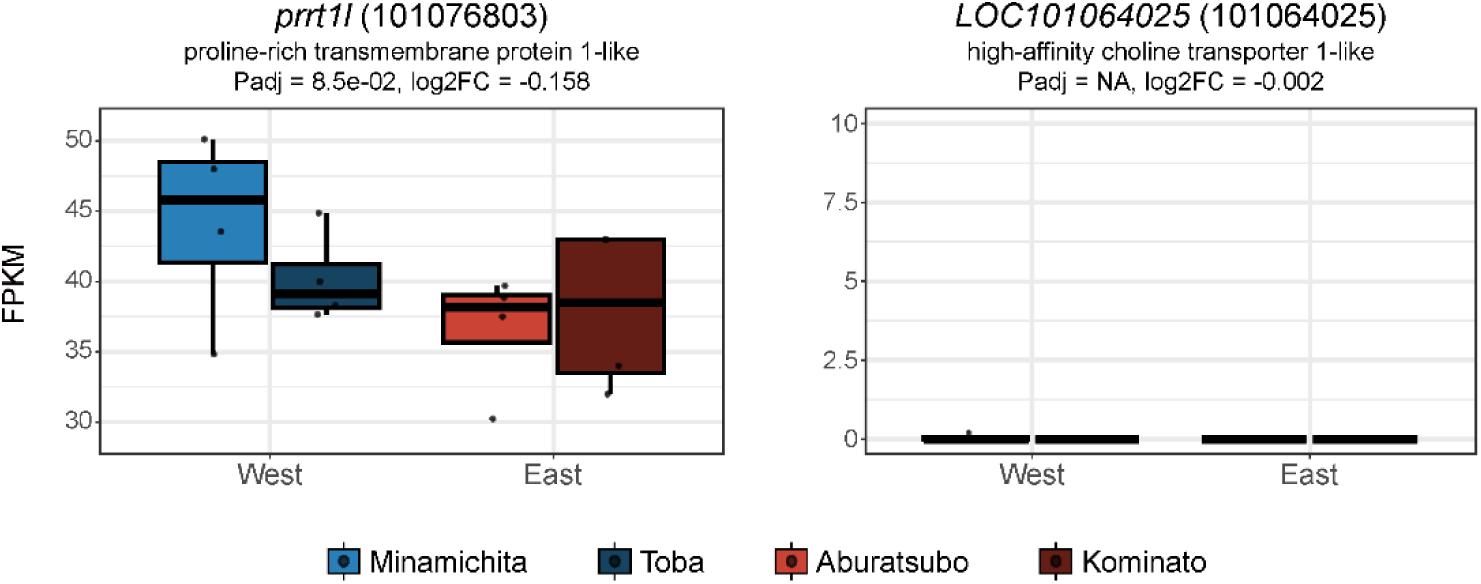
Expression levels of *prrt1l* and *LOC101064025* in the hypothalamus and pituitary, derived from RNA-seq data of western (Minamichita and Toba) and eastern (Aburatsubo and Kominato) populations. *LOC101064025* was hardly expressed in these regions. While *prrt1l* was detected in all samples, no significant differential expression was observed between the two populations.

**Fig. S5.**
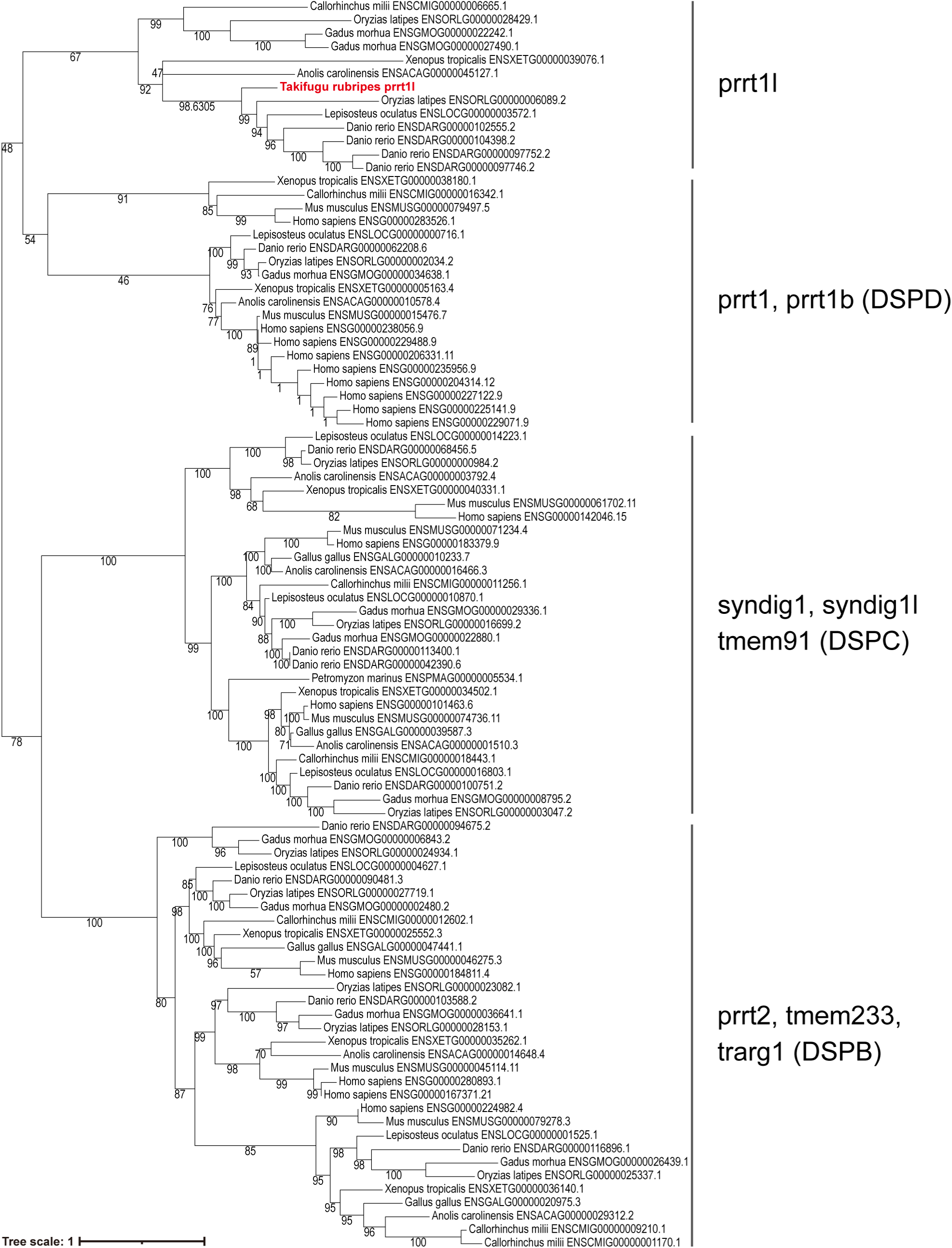
Phylogenetic tree of prrt1l and its relatives generated by SHOOT.bio^51^. The tiger puffer prrt1l is highlighted in red. Representative protein name and, where known, the subfamily names are shown.

## Notes

### Competing Interest Statement

The authors have declared no competing interest.

### Summary of Updates

The mathematical modeling has been added to clarify the mechanistic link between circadian period and the phase of the semilunar rhythm.

