## Supplementary Figures for "Population genomics reveal genetic variants associated with lunar-regulated spawning time in grass puffer"

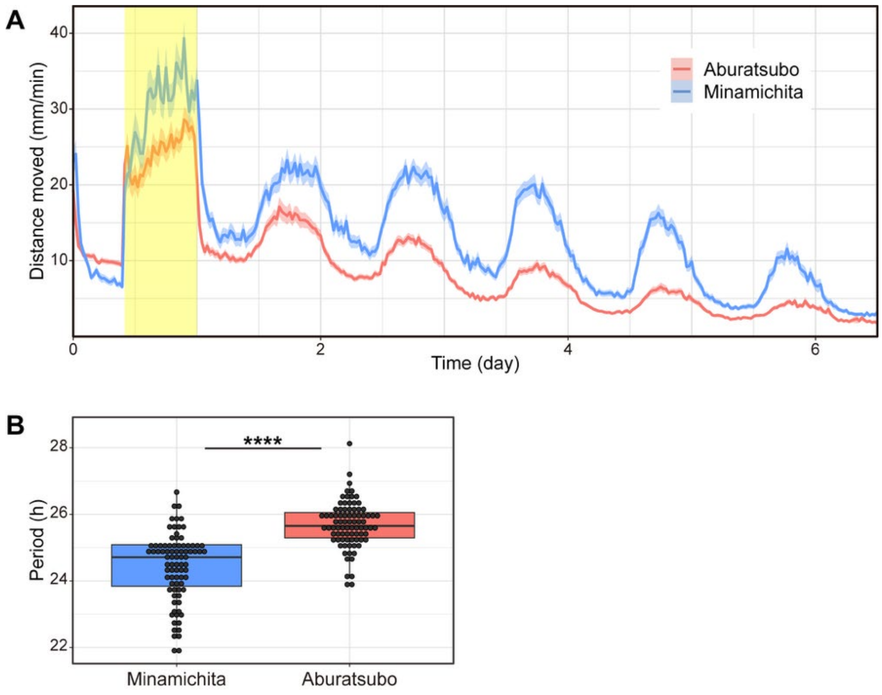


Fig. S1. Shorter free-running period in Minamichita than Aburatsubo. (A) Temporal changes in the average activity levels of larvae in each population analyzed in summer 2025. The solid lines represent the average values, and the shaded areas represent the 95% bootstrap confidence interval. The yellow background represents the light period. (B) The free-running period of larvae from two populations calculated using the Lomb-Scargle periodogram (Aburatsubo: $\tau = 25.68 \pm 0.69$ h, n = 83; Minamichita: $\tau = 24.43 \pm 1.02$ h, n = 83; Welch's *t*-test: $t_{(144.95)} = 9.09$, $p < 0.001$).

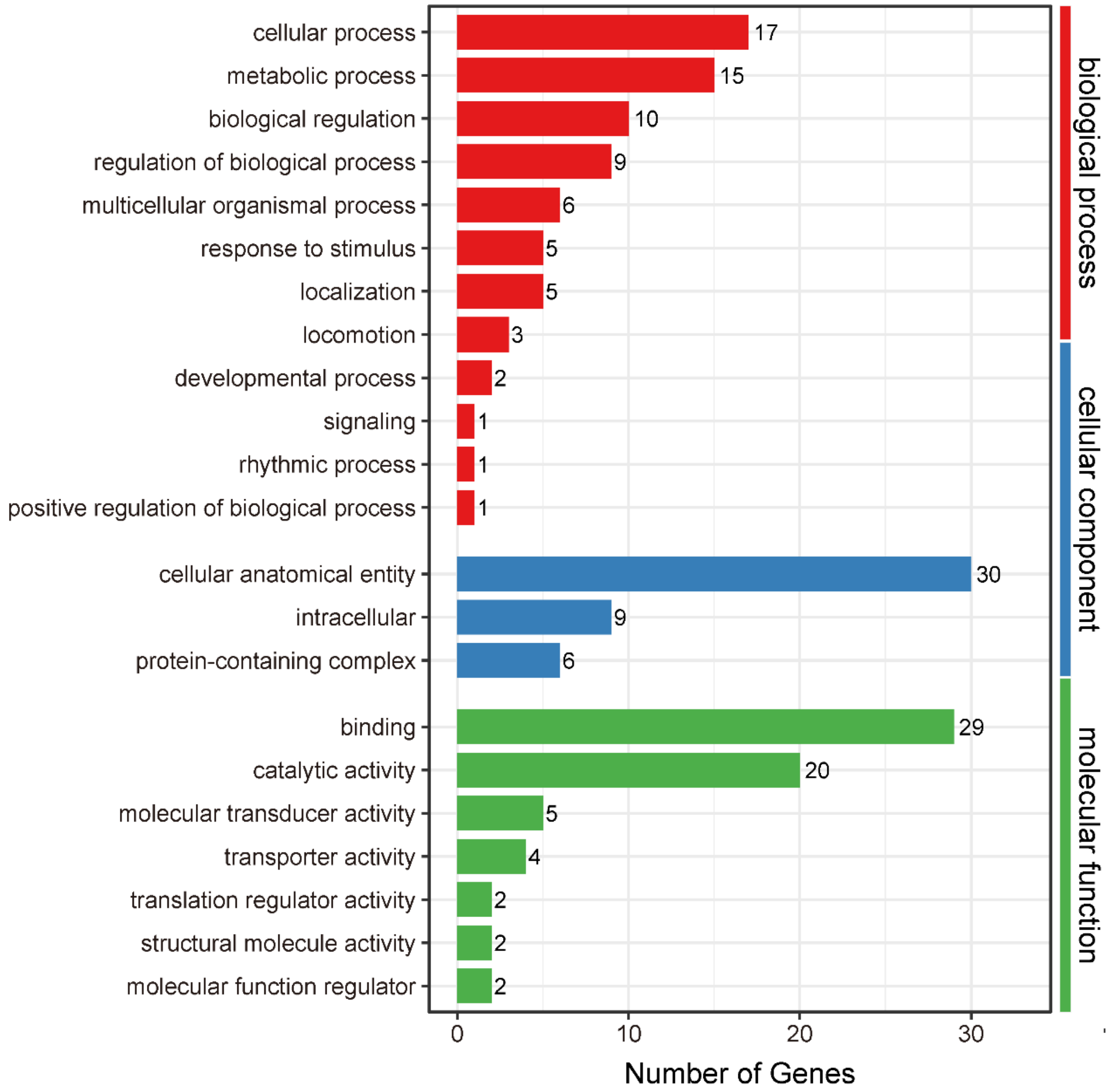


Fig. S2. Gene ontology classification of the 88 DEGs. X axis represents number of DEGs. Y axis represents GO term.

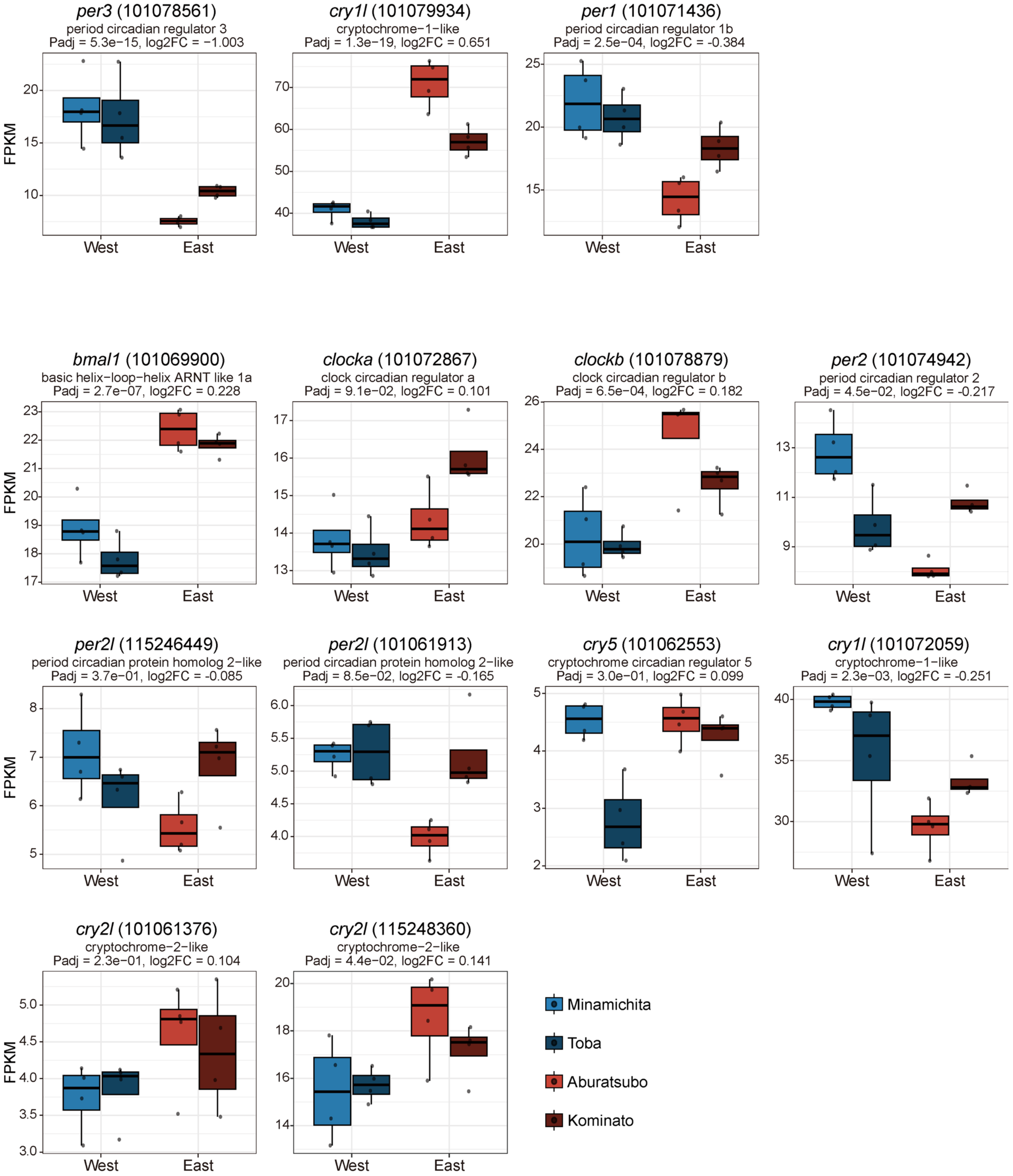


Fig. S3. Expression levels of all core clock genes in RNA-seq data from western (Minamichita and Toba) and eastern (Aburatsubo and Kominato) populations. Clock genes exhibiting different expression patterns between populations are shown in the upper panel and other clock genes are shown in the lower panel. Boxplots show the distribution of expression levels in each population, with black dots representing individual values.

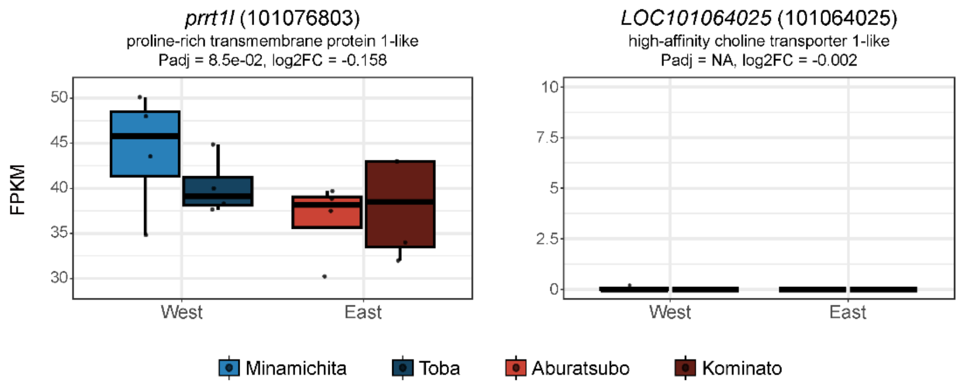


Fig. S4. Expression levels of *prrt1l* and *LOC101064025* in the hypothalamus and pituitary, derived from RNA-seq data of western (Minamichita and Toba) and eastern (Aburatsubo and Kominato) populations. *LOC101064025* was hardly expressed in these regions. While *prrt1l* was detected in all samples, no significant differential expression was observed between the two populations.

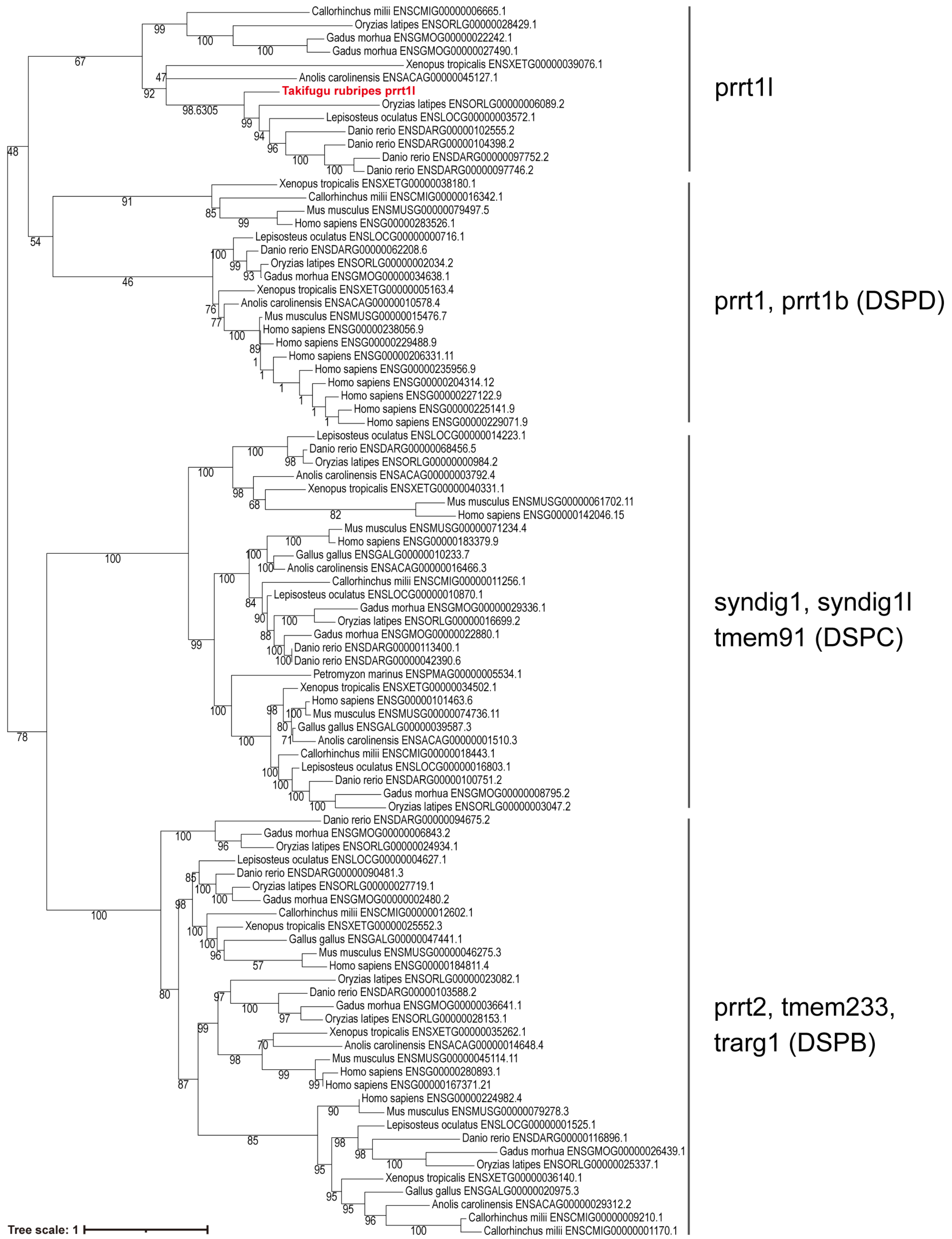


Fig. S5. Phylogenetic tree of prrt1l and its relatives generated by SHOOT.bio[51]. The tiger puffer prrt1l is highlighted in red. Representative protein name and, where known, the subfamily names are shown.
